# The effect of *Drosophila attP40* background on the glomerular organization of Or47b olfactory receptor neurons

**DOI:** 10.1101/2022.06.16.496338

**Authors:** Qichen Duan, Rachel Estrella, Allison Carson, Yang Chen, Pelin C. Volkan

## Abstract

Bacteriophage integrase-directed insertion of transgenic constructs into specific genomic loci has been widely used by *Drosophila* community. The *attP40* landing site located on the second chromosome gained popularity because of its high inducible transgene expression levels. Here, unexpectedly, we found that homozygous *attP40* chromosome disrupts normal glomerular organization of Or47b olfactory receptor neuron (ORN) class in *Drosophila*. This effect is not likely to be caused by the loss of function of *Msp300*, where the *attP40* docking site is inserted. Moreover, the a*ttP40* background seems to genetically interact with the second chromosome *Or47b-GAL4* driver, which results in a similar glomerular defect. Whether the ORN phenotype is caused by the neighboring genes around *Msp300* locus in the presence of *attP40*-based insertions or a second unknown mutation in the *attP40* background remains elusive. Our findings tell a cautionary tale about using this popular transgenic landing site, highlighting the importance of rigorous controls to rule out the *attP40* landing site-associated background effects.

## Introduction

RNA interference (RNAi)-based genetic screens provide scientists with powerful tools to identify genes involved in various biological processes (Housden *et al*. 2017). Binary expression systems, such as the *GAL4/UAS* system, induce the expression of various effectors in the desired cell populations (Brand and Perrimon 1993). In Drosophila carrying transgenes for both cell-type-specific promoter-driven *GAL4* (driver) and *UAS-RNAi*, GAL4 protein binds *UAS* sites and drives *RNAi* expression, disrupting the expression and function of the target gene (Brand and Perrimon 1993). As RNAi-based knockdown methods were becoming popular, efforts were initiated to make transgenic libraries of flies carrying *UAS-RNAi* targeting all the genes in the genome (Dietzl *et al*. 2007; Ni *et al*. 2009; Ni *et al*. 2011; Perkins *et al*. 2015). These genome-wide libraries were then followed by efforts to generate thousands of *GAL4* lines that restrict expression to cellular subpopulations, enabling loss-of-function screens in cells of interest.

Among the RNAi collections, stocks from Transgenic RNAi Project (TRiP) have gained popularity because of their targeted integration of *UAS-RNAi* transgenes into the genome, efficient expression induced by appropriate *GAL4* drivers in different tissues, and high specificity with minimal expected off-target effects (Markstein *et al*. 2008; Ni *et al*. 2008; Perkins *et al*. 2015). To expedite the generation of transgenic libraries, two predetermined chromosomal docking sites were targeted for recombination events that insert *UAS-RNAi* transgenes: *attP40* on the second chromosome and *attP2* on the third chromosome (Markstein *et al*. 2008). With the presence of bacteriophage-originated phiC31 integrase (by co-injection of integrase mRNA or germline-expressing transgenic integrase), the *UAS-RNAi* construct can be inserted into the corresponding docking sites (Groth *et al*. 2004; Ni *et al*. 2008). These two sites, *attP40* and *attP2*, are selected because they exhibit optimal inducible expression levels upon binding with diverse tissue-specific *GAL4* drivers (Markstein *et al*. 2008). Therefore, in addition to the TRiP *UAS-RNAi* library, many other transgenes, including tissue-specific drivers (*GAL4, QF, LexA*) and *UAS/QUAS/LexAop*-*effectors/reporters* are also routinely integrated into these two landing sites (Zirin *et al*. 2020).

Given the widespread use of transgenic flies with *attP40* and *attP2* backbones, and the lesson learned from another popular *UAS-RNAi* collection with reported non-specific effects due to transgenic docking sites (Green *et al*. 2014; Vissers *et al*. 2016), we must be more cognizant of potential phenotypic influences from these genetic backgrounds. Both *attP40* and *attP2* docking sites are in chromosomal regions populated by many genes. These sites, like any insertion into the genome, can disrupt function of nearby genes. More specifically, the *attP40* site is located within one of the large introns of *Msp300* gene while *attP2* site is inserted in the 5’ untranslated region (UTR) of *Mocs1* gene (Larkin *et al*. 2020). Both Msp300 and Mocs1 have critical biological roles. Specifically, Msp300 is the *Drosophila melanogaster* orthologue of mammalian Nesprins, which organize postsynaptic cytoskeleton scaffold and are required for stabilization of new synapses (Elhanany-Tamir *et al*. 2012; Morel *et al*. 2014; Titlow *et al*. 2020; Zheng *et al*. 2020). Mocs1 is involved in Mo-molybdopterin cofactor biosynthetic process and inter-male aggressive behaviors (Gaudet *et al*. 2011; Ramin *et al*. 2019). It is unclear how the insertion of various transgenic constructs into *attP40* and *attP2* docking sites would affect the function of these host genes which may further result in phenotypic defects.

Indeed, recent studies have raised issues related to landing site-associated effects. For example, van der Graaf et al. showed flies bearing two copies of *attP40*-derived insertions also show decreased *Msp300* transcript levels (van der Graaf *et al*. 2022). In addition, this study also reported defects in muscle nuclei spacing in larval stages in the *attP40* homozygous background, which phenocopies *Msp300* mutants (van der Graaf *et al*. 2022). These results suggest that the *attP40* docking site and *attP40*-based transgenes are insertional mutations of *Msp300* gene (van der Graaf *et al*. 2022). Another study reported that *attP40* flies show resistance to cisplatin-induced neuronal damage, compared to the *attP2* background (Groen *et al*. 2022). This study tied the effect to the reduced *ND-13A* (NADH dehydrogenase 13 kDA subunit, a component of mitochondrial complex I) expression in *attP40* homozygous flies (Groen *et al*. 2022). It is noteworthy that *ND-13A* flanks the 5’ UTR of *Msp300* and is downstream of *attP40* docking site. Together, these results imply the integration of *attP40* docking site significantly changes the local transcriptional state and interferes with the transcription of surrounding genes.

During a GAL4-driven *UAS-RNAi* screen for olfactory neuron axon organization, we observed an axon terminal phenotype that is associated with the *attP40* background. The phenotype occurs in the flies homozygous for the *attP40* docking site alone or with various transgenic insertions, independent of the identity of the transgene. Notably, the phenotype observed in the *attP40* background appears to be recessive but is independent of the *Msp300* function, possibly implicating other *attP40* background mutations nearby or in other locations on the second chromosome. Though the nature of the mutation is unclear, the background effects should be mitigated by designing more rigorous controls to interpret phenotypic data obtained using reagents in concert with the *attP40* background.

## Results

We used the *Drosophila* olfactory receptor neurons (ORNs) as a model to understand the molecular mechanisms underlying neuronal circuit assembly. In *Drosophila*, each class of ORNs expresses a unique olfactory receptor (*Or*) gene, and ORN axons target to the brain antennal lobe within class-specific and uniquely positioned synaptic units called glomeruli (Hong and Luo 2014; Barish and Volkan 2015). To identify the molecular players contributing to the glomerular organization of the ORNs, we genetically screened genes encoding cell adhesion molecules whose expression levels increase over pupal development in the antennae (Barish *et al*. 2018). Among these, *beat* and *side* gene families drew our attention because they encode the Ig superfamily proteins, form a heterophilic interacting protein network, and have been previously revealed to be involved in neuronal adhesion (Fambrough and Goodman 1996; Pipes *et al*. 2001; Sink *et al*. 2001; DE Jong *et al*. 2005; Siebert *et al*. 2009; ÖZkan *et al*. 2013; Li *et al*. 2017; Kinold *et al*. 2021). We obtained a collection of transgenic *UAS-RNAi* lines from TriP library deposited at the Bloomington Drosophila Stock Center (BDSC) and crossed these lines with an established recombinant chromosome containing four different *Or* promoter-driven *GAL4* transgenes (*Or47a-GAL4, Or47b-GAL4, Or23a-GAL4, Gr21a-GAL4, 4xOr-GAL4* for short, Figure 1A-D) together with a *UAS-Syt*.*GFP* reporter. We examined the knockdown effect of candidate genes on axonal targeting of these four ORN classes. The parent flies with a single copy of the *GAL4* drivers showed wild type glomerular organization (Figure 1A). As an additional control, we also crossed *GAL4* driver lines to flies expressing the RNAi against a red fluorescent protein (RFP) mCherry, which also exhibited no apparent defect in glomerular organization (Figure 1A).

**Figure 1.**
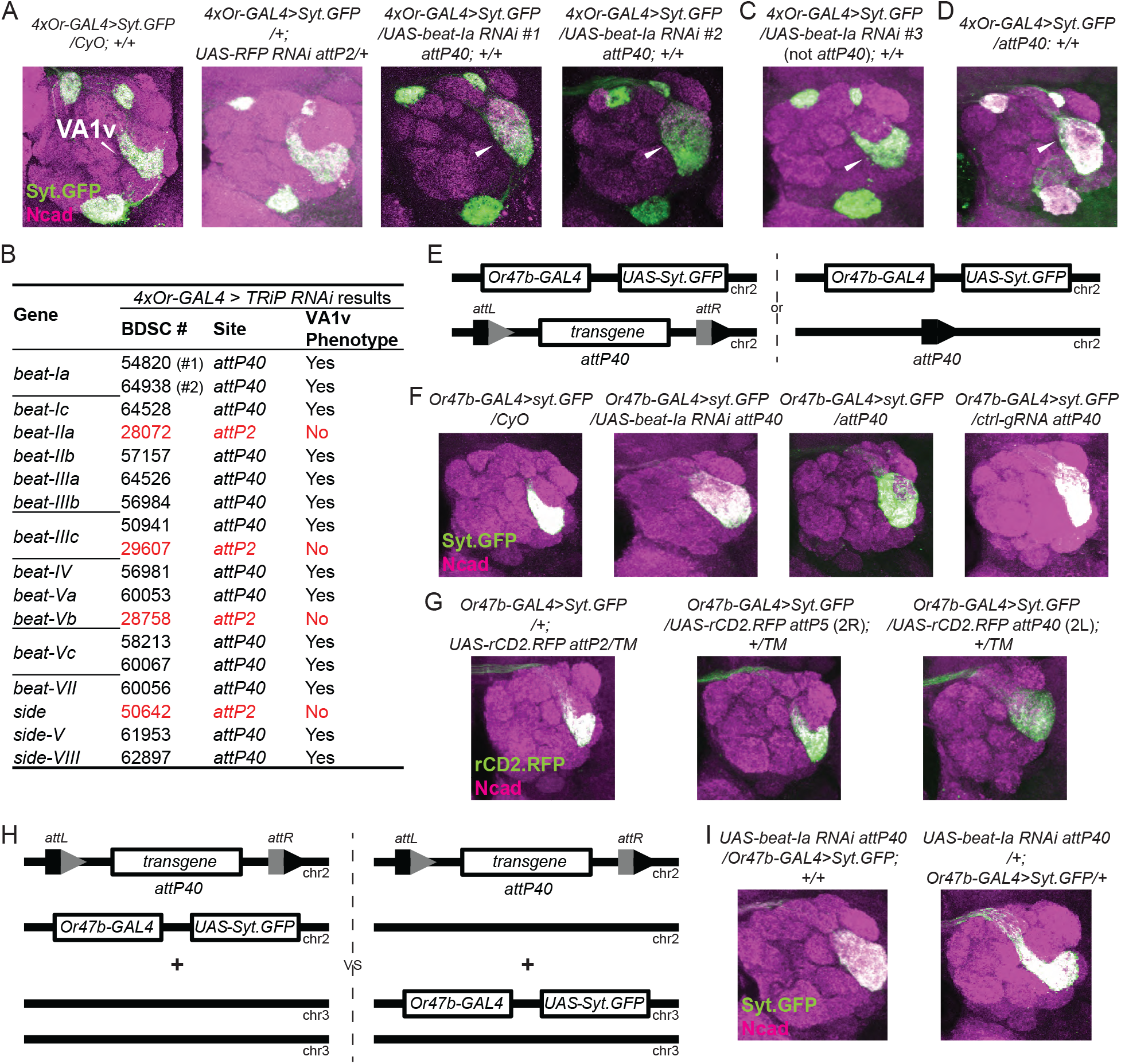
Genetic interactions between *attP40* and *Or47b-GAL4* backgrounds on the second chromosome disrupt the glomerular organization of Or47b ORNs in the antennal lobes. **(A, C & D)** Confocal images of representative brains from a genetic screen to identify adhesion molecules involved in the glomerular organization of the Drosophila olfactory receptor neuron axon terminals. We crossed a second chromosome containing four different Drosophila olfactory receptor promoter-driven *GAL4s* (*Or47a-GAL4, Or47b-GAL4, Or23a-GAL4, Gr21a-GAL4*) together with a *UAS-Syt*.*GFP* reporter (*4xOr-GAL4>Syt*.*GFP*) to the indicated *UAS-RNAi* lines or *attP40* background flies. The parental driver chromosome over the *CyO* balancer was used as a no-RNAi control. The invading Or47b ORN axons are denoted with white arrowheads. **(B)** Summary of the phenotypical results from the genetic screen focusing on *beat/side* gene families. The Bloomington stock number and the transgenic docking site of each line are also listed. **(E, F)** Schematic in **(E)** shows the genotype of animals used in **(F)**, where each fly has one copy of the second chromosome carrying an *Or47b-GAL4* driver and a *UAS-Syt*.*GFP* reporter, and one copy of the indicated second chromosome, either a *CyO* balancer or *attP40* docking site derivatives. *attL* and *attR* sites are generated as a result of transgene integration into *attP40* docking site. Confocal images of representative brains are shown in **(F). (G)** Confocal brain images of the indicated genotypes. **(H, I)** Schematic in **(H)** shows the genotype of animals used in **(I)**, where each fly has one copy of the second chromosome *UAS-beat-Ia RNAi* transgene inserted into the *attP40* docking site, with one copy of *Or47b-GAL4 UAS-Syt*.*GFP*, either on the second or third chromosome. Confocal images of representative brains are shown in **(I)**. 10-25 brains were examined in each genotype and the phenotypical penetrance is close to 100% in each *attP40*-derived group.

From the screen, we found a strikingly recurrent phenotype, where the axon terminals of Or47b ORNs invade the neighboring region, leading to an expanded round VA1v glomerulus in contrast to the crescent shape in control brains (Figure 1A). This phenotype was observed in two independent RNAi lines targeting the same gene, for example, *beat-Ia* (Figure 1A). However, screening a list of *beat* and *side* family members revealed a pattern for the phenotype, which only correlated with the second chromosome *UAS-RNAi* transgenes, independent of the gene identity. Figure 1B summarizes the screening results from *beat/side* gene families. All the RNAi lines inserted at the second chromosome *attP40* site yielded the expanded VA1v glomerulus phenotype, whereas none of the RNAi lines inserted at the third chromosome *attP2* site showed this defect. Notably, there is one gene, *beat-IIIc*, with one *attP40*-derived RNAi line and one *attP2*-derived RNAi line. Only the *attP40 UAS-RNAi* insertion gave rise to the phenotype (Figure 1B). The same phenotype was also observed with randomly selected TRiP *UAS-RNAi* lines inserted at the *attP40* site targeting genes without known roles in ORN development (Figure S1A,B). To test whether this phenotype is caused by specific effects of RNAi-mediated gene knockdown or simply by the presence of *attP40*-derived insertions, we first crossed the same *Or47b-GAL4* driver line to a third *UAS-RNAi* line from Vienna Drosophila Resource Center (VDRC) targeting *beat-Ia*, which was generated by random P-element-mediated insertions (Dietzl *et al*. 2007). This non-*attP40 UAS-RNAi* line could not reproduce the phenotype obtained by the *attP40*-derived *UAS-RNAi* from the TRiP collection (Figure 1C). In addition, crossing the driver line to an empty *attP40* site without any transgenes led to the same glomerular expansion phenotype (Figure 1D). These results suggest that the Or47b ORN-specific VA1v glomerular defect is independent of the RNAi-based knockdown of the genes examined but caused by an effect from the *attP40*-derived chromosome.

Since we repeatedly obtained the VA1v glomerular phenotype with the second chromosome *Or47b-GAL4*-driven *UAS-RNAi*, we also tested if crossing flies carrying the same *Or47b-GAL4* transgene to various *attP40* derivatives could result in the same phenotype (Figure 1E). Compared with the no *attP40* control (over a *CyO* balancer chromosome), the *attP40* landing site with and without *UAS-RNAi* insertion, or a ubiquitous promoter-driven gRNA targeting the *QUAS* sequence (control gRNA) all produced the same VA1v glomerular defect when crossed to the second chromosome *Or47b-GAL4*-driven *UAS-Syt*.*GFP* (Figure 1F). We also crossed *Or47b-GAL4 UAS-Syt*.*GFP* chromosome to three lines carrying *UAS-rCD2*.*RFP* transgenic insertion at three different chromosomal locations, *attP2* (on chr3), *attP5* (on chr2R), and *attP40* (on chr2L). Only *Or47b-GAL4* over *UAS-rCD2*.*RFP* insertion at *attP40* resulted in glomerular expansion phenotype while the insertions at *attP2 and attP5* appeared wild type. These results again suggest that the VA1v glomerular defect is uniquely linked to the *attP40*-associated insertions and is independent of the transgene or other attP landing sites.

The glomerular organization defect could be caused by simply the presence of *attP40* insertion or the genetic interaction between the *attP40* background and the chromosome carrying the reporter transgene. To distinguish between these possibilities, we examined the animals carrying a single copy of *attP40* insertion and a different *Or47b-GAL4 UAS-Syt*.*GFP* reporters on the third chromosome. In these animals, VA1v glomerulus appeared normal (Figure 1H,I). The observation that a single copy of *attP40* is not sufficient to produce a glomerular phenotype indicates that the *attP40* effects on VA1v glomerulus are not dominant. Rather, they point to a combinatorial effect of the second chromosome with *Or47b-GAL4 UAS-Syt*.*GFP* over the chromosome with the empty or transgene-carrying *attP40* docking site on the glomerular phenotype (Figure 1H,I). Thus, we conclude that the *attP40* chromosome genetically interacts with the second chromosome reporters to disrupt VA1v glomerular organization.

As the *attP40* effect appears to be recessive, we next examined if animals homozygous for the *attP40* sites display any VA1v glomerular defects. To bypass the glomerular defects arising from the genetic interactions between *attP40* and the second chromosome reporters, we used the third chromosome *Or47b-GAL4 UAS-Syt*.*GFP* reporter to visualize VA1v glomerulus. Surprisingly, homozygous *attP40* derivatives or *attP40* empty docking site alone produced strong axon terminal defects (Figure 2A-F). In contrast, flies heterozygous for the *attP40* site with or without transgenes inserted appeared wild type (Figure 2A-F). Most of the brains homozygous for the *attP40* site with or without insertions displayed a dorsally positioned VA1v glomerulus (Figure 2B,E, middle panels; Figure 2C,F), whereas a small proportion also exhibited an expanded glomerulus (Figure 2B,E, right panels; Figure 2C,F). Given that the *attP40* site is located within an intron of *Msp300* gene, we posited that it likely disrupts *Msp300* function. *Msp300* encodes a Nesprin-like protein, which is required for proper positioning of muscle nuclei and neuromuscular junction formation (Elhanany-Tamir *et al*. 2012; Morel *et al*. 2014). Single-cell RNA-seq datasets from ORNs also show broad expression of *Msp300* across ORN classes (Li *et al*. 2022). We thus tested if the VA1v glomerular defect is caused by the loss of *Msp300* function. We analyzed transheterozygotes of empty *attP40* docking site over other mutant alleles of *Msp300*, such as *Msp300*^*ΔKASH*^ (which lacks the KASH domain (Xie and Fischer 2008; Elhanany-Tamir *et al*. 2012)), *Msp300*^*MI00111*^, *Msp300*^*MI01145*^ (two MIMIC-based alleles predicted to disrupt most splice isoforms of *Msp300* transcripts (Venken *et al*. 2011)), and *Msp300*^*KG03631*^ (a P-element-based insertion which is close to *attP40* landing site (Bellen *et al*. 2004)) (Figure 2G). However, none of these genetic combinations recapitulated VA1v glomerular phenotype (Figure 2H). This indicates that the VA1v glomerular defect is independent of the *Msp300* function and is likely caused by other genes nearby affected by the *attP40* insertion or a second recessive mutation linked to the *attP40* docking site.

**Figure 2.**
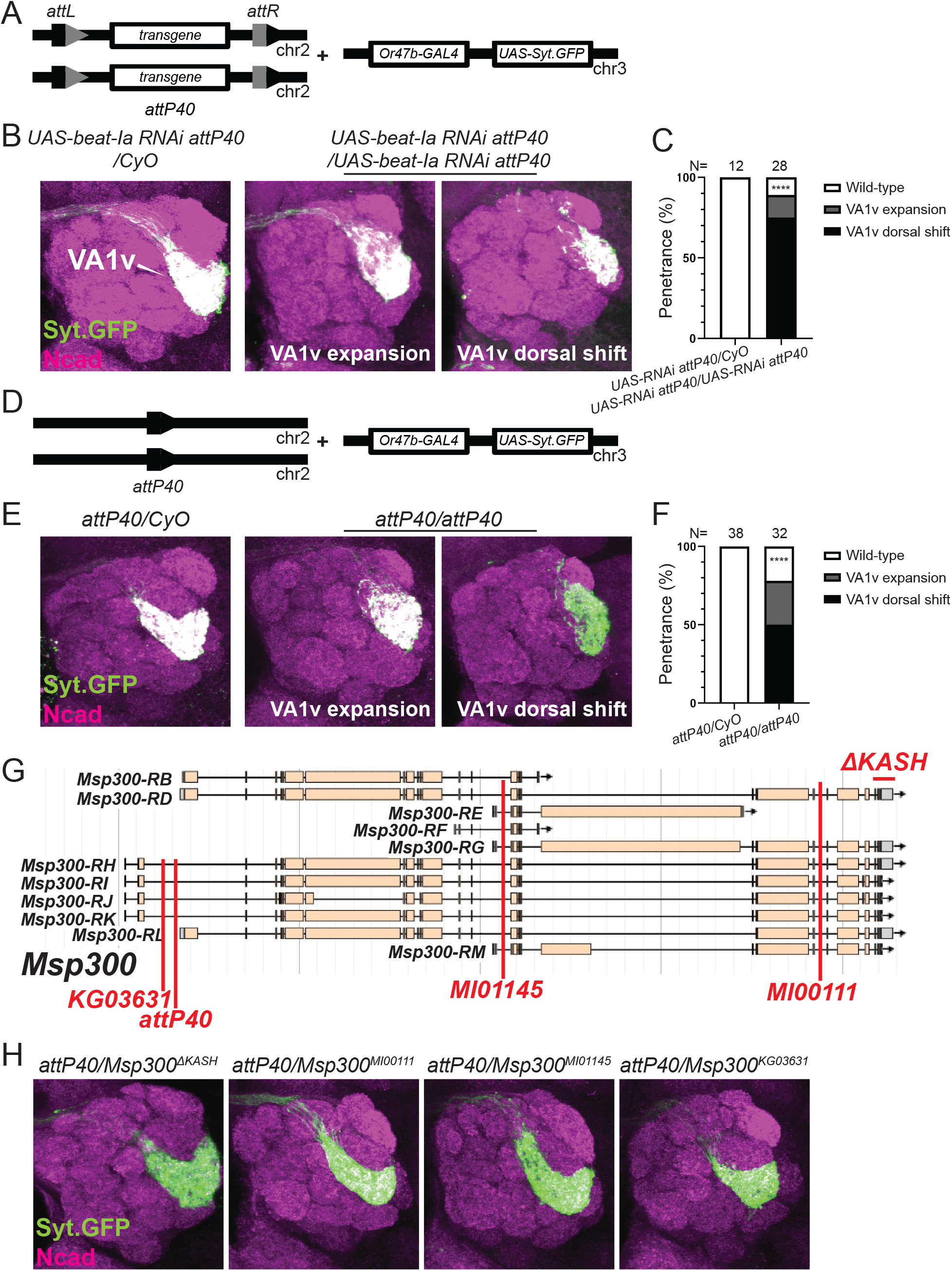
Homozygous *attP40* chromosome affects glomerular organization of Or47b ORNs independent of the *Msp300* function. **(A-C)** Schematic in **(A)** shows the genotype of animals used in **(B)**, where each fly has one or two copies of the second chromosome *UAS-beat-Ia RNAi* transgene inserted at the *attP40* docking site, with the third chromosome *Or47b-GAL4 UAS-Syt*.*GFP* transgenes. Confocal images of representative brains are shown in **(B)**. The percentage of the phenotypes is shown in **(C)**. ****, p<0.0001 after Fisher’s exact test. **(D-F)** Schematic in **(D)** shows the genotype of animals used in **(E)**, where each fly has one or two copies of the second chromosome empty *attP40* docking site, with the third chromosome *Or47b-GAL4 UAS-Syt*.*GFP* transgenes. Confocal images *of* representative brains are shown in **(E)**. The percentage of the phenotypes is shown in **(F)**. ****, p<0.0001 after Fisher’s exact test. N in **(C)** and **(F)** denotes the antennal lobes examined. **(G)** Schematic showing the *Msp300* genomic locus, the *attP40* docking site, three insertional *Msp300* mutations (*Msp300*^*MI00111*^, *Msp300*^*MI01145*^, *Msp300*^*KG03631*^), and one deletion allele (*Msp300*^*ΔKASH*^), each denoted with red lines. **(H)** Confocal images of representative brains of the indicated transheterozygous animals, with the *attP40* docking site over the indicated *Msp300* alleles. N = 11, 8, 4, 12 brains in each genotype group, from left to right.

We next sought to figure out the molecular basis of the genetic interaction between that specific *Or47b-GAL4*-bearing chromosome and *attP40* chromosome. Flies transheterozygous for *attP40* (empty or with insertions) over *4xOr-GAL4 UAS-Syt*.*GFP* or *Or47b-GAL4 UAS-Syt*.*GFP* robustly exhibit VA1v glomerular phenotype. We infer that the putative genetic lesion is directly caused by or genetically linked to the *Or47b-GAL4* transgene for three reasons: 1) a farther second site mutation would likely be lost during meiotic recombination events to generate these stocks; 2) this *Or47b-GAL4* recombined with other *UAS-reporters, UAS-mCD8*.*GFP* or *UAS-RFP*, over the *attP40* derivatives exhibited the same phenotype (Figure S2); and 3) crossing the second chromosome *Or47b-GAL4* transgene alone to the *UAS-rCD2*.*RFP* reporter inserted at *attP40* also reproduced the expanding glomerulus (Figure 3A), which rules out the confounding effect from *UAS-Syt*.*GFP* transgene. Indeed, two copies of this *Or47b-GAL4* chromosome also results in similar glomerular phenotypes (Figure 3A). *Or47b-GAL4* transgene was generated by P-element-mediated genomic integration (Figure 3B) and the exact site of the insertion was not mapped (Vosshall *et al*. 2000). We used inverse PCR (Huang *et al*. 2009) to identify the insertion site and determine the gene whose function is potentially disrupted (Figure 3C). We successfully recovered a piece of genomic sequence immediately flanking the 3’ end of the inserted P-element, which is within the first intron of *Bacc* gene (chr2L:2753160, Figure 3D,E) and also a P-element insertion-enriched region. We performed genomic PCR to further validate the *Bacc* intronic insertion from both 3’ and 5’ ends of the P-element using different primer pairs targeting the P-element ends and flanking *Bacc* sequences (Figure 3E). We amplified the desired DNA fragments from *Or47b-GAL4*-bearing flies but not from *w*^*1118*^ flies (Figure 3E,F). In addition, primer pairs targeting only *Bacc* sequences flanking the insertion amplified the expected fragment only from *w*^*1118*^ files but not from *Or47b-GAL4* flies (Figure 3E,F). We next tested if this intronic insertion affects *Bacc* transcriptional levels using Quantitative Reverse Transcription-PCR (qRT-PCR). We extracted mRNA from whole heads of the adult *Or47b-GAL4* homozygotes, as well as homozygous *attP2, attP40*, and *w*^*1118*^ adults. qRT-PCR results showed that *Bacc* transcripts normalized to the housekeeping gene *RpL13A* decrease by ∼two fold in flies homozygous for *Or47b-GAL4* but are not significantly altered in *attP2* or *attP40* animals compared to *w*^*1118*^ controls (Figure 3G). In contrast, other housekeeping genes *Act5C* and *Tbp* remain unchanged across these genotypes (Figure 3G). These results indicate that: 1) second chromosome *Or47b-GAL4* transgene insertion disrupts *Bacc* gene function; 2) homozygous and transheterozygous combinations of *Or47b-GAL4* and *attP40* backgrounds likely utilize distinct mechanisms to disrupt VA1v glomerular organization.

**Figure 3.**
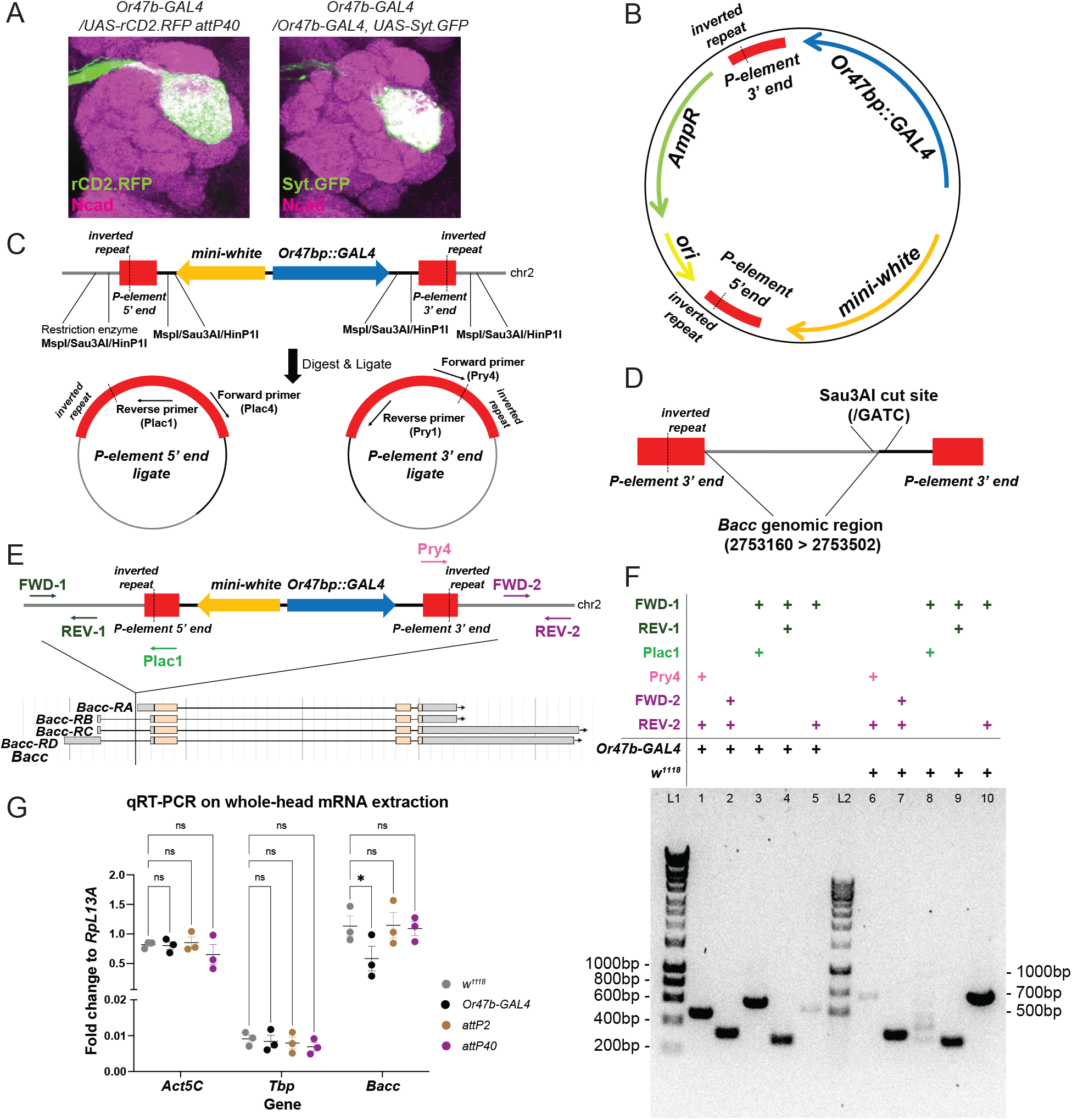
Identification of the *Or47b-GAL4* transgene insertion site. **(A)** Confocal images of the indicated genotypes suggesting the second chromosome *Or47b-GAL4* transgene is accounting for the glomerular expansion phenotype. N = 16 brains in each genotype and the phenotypical penetrance is 100%. **(B)** The structure of the P-element vector containing *Or47b-GAL4* transgene (*Or47b* promoter fused with GAL4 coding sequence, denoted as *Or47bp::GAL4* in this and other panels of this figure) and a *mini-white* selectable marker. **(C)** Schematic illustrating the recovery of genomic sequences flanking P-element insertion by inverse PCR. **(D)** Schematic showing the sequencing results from one ligated genomic DNA template digested by Sau3AI restriction enzyme. A region within *Bacc* gene was recovered as the immediate sequencing flanking the 3’ end of the inserted P-element containing *Or47b-GAL4* transgene. **(E)** Schematic showing the P-element insertion site within the first intron of *Bacc* gene. The arrows indicate the primers used in a PCR assay to validate this insertion. **(F)** DNA gel showing the amplicons by each indicated primer pairs from *Or47b-GAL4* transgenic flies or *w*^*1118*^ control flies. Primer pairs FWD-1/REV-1 and FWD-2/REV-2 can amplify the specific fragments from both *Or47b-GAL4* transgenic flies and *w*^*1118*^ control flies as expected because the corresponding sequences exist in both. Primer pairs FWD-1/Plac1 and Pry4/REV-2 only amplify specific products from *Or47b-GAL4* transgenic flies while fail to work from *w*^*1118*^ control flies as one primer in each pair targets the sequence of P-element which is not present in *w*^*1118*^ flies. Primer pair FWD-1/REV-2 works in *w*^*1118*^ control flies but fails in *Or47b-GAL4* flies as the expected amplicon is too long to be amplified in transgenic animals because of the integration of large P-element sequence. **(G)** qPCR results comparing the indicated gene expression levels normalized by *RpL13A* transcripts across different genotypes. ns, not significant. *, p-value < 0.05 after two-way ANOVA followed by multiple comparison test. Error bars indicate mean ± SEM. N = 3 biological replicates per genotype. Each biological replicate contain 14∼20 fly heads from equal number of males and females.

## Discussion

Here, we found that homozygous *attP40* chromosome leads to defective glomerular organization of ORNs. This defect is likely not caused by the loss of *Msp300* function, where the *attP40* site is inserted. Moreover, the a*ttP40* chromosome genetically interacts with a second chromosome carrying the *Or47b-GAL4* transgene, resulting in a similar ORN axon terminal defect. Though the exact genetic reasons and molecular mechanisms are unknown, our finding raises the critical issue with using this popular transgene landing site. Rigorous controls are needed to rule out the *attP40*-associated background effects, as discussed below.

### The genetics underlying the Or47b ORN phenotypes

A recent study reported that flies homozygous for the *attP40*-derived insertions had 50% reduction in *Msp300* transcript levels and phenocopied the defects in larval muscle nuclei clustering in *Msp300* mutants (van der Graaf *et al*. 2022). As homozygotes of the *attP40* chromosome are defective in Or47b ORN axon terminal organization, we hypothesized that the *attP40*-affected *Msp300* gene is responsible for the defect. However, this is not the case as *attP40* over various *Msp300* mutations appeared phenotypically wild type, suggesting the *attP40* chromosome may carry an unannotated mutation responsible for *Msp300*-independent ORN glomerular disorganization.

The *attP40* docking site with or without transgene insertions may also disrupt other genes in the vicinity of *Msp300*. For example, in addition to *Msp300, attP40* docking site is flanked on the opposing side by *ND-13A*, which encodes a component of the mitochondria electron transport chain complex I. Thus, the *attP40* docking site alone or with transgene insertions may lead to a variety of phenotypes as a result of disrupted *ND-13A*. Indeed, Groen et al. reported that *attP40* flies exhibit resistance to cisplatin-induced neuronal damage mediated by the reduced expression of *ND-13A* (Groen *et al*. 2022). Whether the glomerular defect is dependent on the *ND-13A* function is beyond the scope of this paper but needs to be tested in the future studies.

Surprisingly, we found transheterozygous animals with an *attP40* chromosome over the second chromosome *Or47b-GAL4* transgene produced similar but not identical glomerular abnormalities to *attP40* homozygotes. Additionally, *Or47b-GAL4* homozygotes exhibit comparable phenotypes with *Or47b-GAL4/attP40* transheterozygotes. These suggest several possible underlying genetic mechanisms. 1) *Or47b-GAL4* and *attP40* backgrounds harbor common mutations; 2) *Or47b-GAL4* and *attP40* backgrounds possess completely separate genetic lesions that genetically interact. The genetic interaction model is favored due to qualitatively distinguishable phenotypes between *Or47b-GAL4/attP40* animals and *attP40/attP40* ones. Furthermore, *Or47b-GAL4* transgene is inserted into an intron of *Bacc* gene, which encodes a tyramine-dependent nuclear regulator (Chen *et al*. 2013), reducing its expression levels by about 50% in homozygotes. No change in transcript levels were observed in *attP40* homozygotes. *Bacc* mRNAs are abundant in brain tissues, comparable to housekeeping genes *Act5C* and *RpL13A* (Figure 3G). Our results imply its potential novel role in ORN axon pathfinding or glomerular patterning. Future functional studies will determine whether the disruption of *Bacc* expression is causative to the VA1v glomerular phenotype and the mechanisms by which *Bacc* mutations and their genetic interactors in the *attP40* background result in glomerular defects.

### Unique genetic sensitivity of VA1v glomerulus architecture

One of the most peculiar observations from our study is that Or47b ORNs seem to be particularly sensitive to changes in genetic background. In fact, VA1v glomerular disruptions are not only restricted to the *attP40* background, but can be seen in many other mutants with effects on ORN axon and synapse organization in the antennal lobes (Ang *et al*. 2003; Yao *et al*. 2007; Hong *et al*. 2012; Li *et al*. 2013; Hueston *et al*. 2016; Wu *et al*. 2017; Xie *et al*. 2019; Hing *et al*. 2020). In addition to the variability of VA1v glomerular architecture to genetic background effects, pheromone sensing Or47b ORNs and the trichoid at4 sensillum that houses Or47b, Or88a, and Or65a/b/c ORNs are developmentally special. At4 sensillum appears to be a developmentally default state for all trichoid sensilla. For example, loss of transcription factor Rn function, normally expressed in at1 and at3 ORNs, leads to a loss of at1 and at3 sensilla identity, and their conversion to at4 sensillum identity (Li *et al*. 2013; Li *et al*. 2015; Li *et al*. 2016). Similarly, Or47b ORNs in at4 sensillum appear to have a default identity, as mutants in *Alh*, a chromatin factor, result in the conversion of Or88a and Or65a ORNs to Or47b ORN fate (Hueston *et al*. 2016). In addition to these findings that point to a developmentally special state for pheromone sensing Or47b ORNs or “at” sensilla, the glomeruli targeted by the trichoid ORNs are morphologically plastic. In Drosophila, they are sexually dimorphic, appearing larger in males (Stockinger *et al*. 2005). In insects such as moths trichoid glomeruli can form separate macro-glomerular complex outside the antennal lobe of male brains (Berg *et al*. 1998). Given the developmental plasticity of trichoid pheromone sensing ORNs and the developmental ground state of at4 sensilla, Or47b developmental trajectory might be particularly sensitive to genetic background effects to accommodate adaptive developmental, behavioral and evolutionary processes. On the other hand, we only examined Or47b VA1v glomerulus in the *attP40* background, and *attP40* homozygotes possibly display structural defects in other ORN classes and their glomeruli. Future studies will help identify these phenotypes and the genetic lesions leading to *attP40*-associated phenotypes.

### Addressing genetic background issues when using genetic reagents

To summarize, we found unexpected background effects of the *Drosophila attP40* landing site on the ORN glomerular organization. In parallel with other recent studies reporting other phenotypes arising from the *attP40* background, ranging from muscle development to neuronal stress responses, such background effects should be seriously considered in using *attP40*-derived flies. It is recommended to avoid using homozygotes/double-copies of the *attP40*-based insertions. Researchers should also be aware of the potential genetic interactions between the *attP40*-bearing chromosome and the other homologous second chromosomes even if it doesn’t contain any *attP40* derivatives. Appropriate controls should be applied to override these caveats. For example, when working with *GAL4/UAS-effector* binary system, it is better to use a GAL4-driven *UAS-neutral effector* (such as *UAS-RNAi* against neutral or non-fly genes inserted at the same docking site) as a negative control, rather than the widespread use of *GAL4* alone or *UAS-effector* alone controls. Transgenic rescue of RNAi-based gene knockdowns is not feasible due to targeting of rescue transgenes by the RNAi. Thus, use of full animal mutants or MARCM based clonal mutant analysis should be coupled with RNAi-based phenotypic analyses. Though the underlying genetic reasons remain elusive, studies demonstrated that the *attP40* landing site on the second chromosome affects the expression of multiple genes (Groen *et al*. 2022; van der Graaf *et al*. 2022). Additional omics-based experiments in the future will be needed to determine all the genetic lesions in *attP40* strains that underly many phenotypic defects observed in this background. These studies will also reveal potential genetic alterations associated with glomerular defects, providing new insights into ORN axon pathfinding and glomerular organization.

## Materials and Methods

### *Drosophila* stocks and genetics

Drosophila were raised in classic molasses media provided by Archon Scientific. For the RNAi screen experiments, flies were raised at 28 °C to maximize the knockdown efficiency. Most of the other crosses were also kept at 28 °C, except for the experiments shown in Figure 1F,G and Figure 3A, which were conducted at room temperature (23 °C). After eclosion, the flies are aged for 5-7 days before dissection. In addition to the *UAS-RNAi* stocks from Bloomington Drosophila Stock Center (listed in Figure 1B), the following stocks are used: *UAS-RFP RNAi attP2* (BDSC# 35785), *UAS-beat-Ia RNAi #3* GD1386 (VDRC# 4544), *UAS-SMC3 RNAi attP2* (BDSC# 60017), *UAS-SMC3 RNAi attP40* (BDSC# 50899), *UAS-vtd RNAi attP2* (BDSC# 36786), *UAS-vtd RNAi attP40* (BDSC# 65229), *attP40* (BDSC# 36304), *attP2* (BDSC# 36303), *ctrl-gRNA attP40* (BDSC# 67539), *UAS-RFP attP40* (BDSC# 32222), *UAS-rCD2*.*RFP attP2* (BDSC# 56179), *UAS-rCD2*.*RFP attP5* (BDSC# 56180), *UAS-rCD2*.*RFP attP40* (BDSC# 56181), *Msp300*^*ΔKASH*^ (BDSC# 26781), *Msp300*^*MI01145*^ (BDSC# 53050), *Msp300*^*MI00111*^ (BDSC# 30623), *Msp300*^*KG03631*^ (BDSC# 13024); *Or47b-GAL4* (chr2, BDSC#9983), *Or47b-GAL4* (chr3, BDSC#9984), *Or43a-GAL4* (chr2), *Or47a-GAL4* (chr2) (Vosshall *et al*. 2000; Fishilevich and Vosshall 2005), and *Gr21a-GAL4* (chr2) (Scott *et al*. 2001)) are gifts from Dr. Leslie Vosshall; *UAS-Syt*.*GFP* (chr2 or chr3), *UAS-mCD8*.*GFP, UAS-RFP* are Volkan lab stocks (Barish *et al*. 2018). The line *Or47b-GAL4, Or47a-GAL4, Or43a-GAL4, Gr21a-GAL4, UAS-Syt*.*GFP/CyO* (short for *4xOr-GAL4>Syt*.*GFP*) was recombined and balanced from the above components.

### Immunocytochemistry

Flies were sacrificed in 70% ethanol. Fly brains were then dissected in PBST buffer (0.2% Triton X-100 in 1X PBS), fixed in 4% paraformaldehyde for 30 mins, followed by washing with PBST for three 10-min cycles. Brains were incubated in the primary antibody mix at 4 °C overnight, followed by three 20-min washes with PBST at room temperature, then incubated in the secondary antibody mix at 4 °C overnight. The brains were washed again by three 20-min wash with PBST before being mounted on the slide for imaging. The blocking was done together with each antibody incubation, with 1% natural goat serum mixed with primary and secondary antibodies, respectively. The following primary antibodies were used: 1:1000 rabbit anti-GFP (Invitrogen), 1:20 rat anti-Ncad (DSHB); the following secondary antibodies were used: 1:1000 Alexa Fluor 488 goat anti-rabbit IgG (Invitrogen), 1:200 Alexa Fluor 647 goat anti-rat IgG (Invitrogen); all antibodies are diluted in PBST.

### Confocal imaging and phenotypic quantification

Confocal imaging was performed by either Olympus Fluoview FV1000 microscope or Zeiss 880 microscope. Brains were imaged across Z-axis from the posterior side to the most anterior side of the antennal lobes, and all confocal sections were overlayed for phenotypical analysis. The same set of imaging parameters was used between experimental and control groups. The phenotype was qualitatively determined by glomerular morphology, i.e., whether Or47b ORN axons appear in the dorsal antennal lobe region, in contrast to the typical V-shaped glomerulus in wild-type controls. The phenotype shown in Figure 1 (glomerular expansion) is largely consistent from brain to brain, while the phenotypes shown in Figure 2B,E exhibit variability, which were categorized into expansion or dorsal shift. The phenotype was quantified by the percentage of antennal lobes exhibiting each defect among all the brains examined in respective groups. P-value was calculated by two-tailed Fisher’s exact test through the built-in functions of GraphPad Prism 9 software.

### Inverse PCR to recover the genomic DNA sequence flanking the *Or47b-GAL4* transgenic insertion

*Or47b-GAL4* transgene was inserted into an unknown region on the second chromosome by P-element-mediated method (Vosshall *et al*. 2000). The P-element structure of *Or47b-GAL4* transgene was shown in the Figure 3B. We used previously described inverse PCR method (Huang *et al*. 2009) to identify the genomic sequence flanking the insertion site. Briefly, genomic DNA was first extracted from 30 *Or47b-GAL4* flies (BDSC# 9983), followed by overnight digestion with any of three restriction enzymes, MspI, HinP1I, or Sau3AI (New England BioLabs). Each digest was then ligated by T4 DNA ligase (New England BioLabs) in larger volume (400 μL, see (Huang *et al*. 2009) for mix details) to promote intramolecular ligation while minimizing intermolecular ligation. Ligation was performed at 16 °C overnight. The unknown flanking sequence was then amplified from the ligated genomic DNA by inverse PCR using forward and reverse primers targeting the 3’- or 5’-end sequences of the P-element (Figure 3C). PCR reaction: 10 μL ligated DNA, 2 μL 5 μM forward and reverse primer mix, 10 μL 5X *myTaq* reaction buffer, 0.5 μL *Taq* DNA polymerase. Thermal cycling program: 3 min initial denaturation at 95 °C + 35 cycles (30 s denaturation at 95 °C, 1 min annealing at 55 °C, 2 min extension at 72 °C) + 10 min final extension at 72 °C. PCR products were then cleaned with Qiagen QIAquick PCR purification kit followed by commercial DNA sequencing service.

A genomic region on the second chromosome (starting from 2L: 2753160) within the first intron of *Bacc* gene was identified (Figure 3D,E) as the immediate sequence flanking the inserted P-element 3’ end. To verify this identified genomic insertion site, a PCR assay was designed to amplify the genomic regions from *Or47b-GAL4* transgenic flies and *w*^*1118*^ control flies respectively with different primer pairs (Figure 3E,F). PCR mix recipe and thermal cycling programs were the same as abovementioned. Amplicons were also purified and sequenced for validation.

### Quantitative Reverse Transcription-PCR

To measure whether the insertion of *Or47b-GAL4* transgene into *Bacc* gene locus affects its expression, we used the quantitative Reverse Transcription-PCR (qRT-PCR) to extract and quantify the mRNA levels from whole heads of fruit flies. RNA extraction, cDNA preparation, and qPCR protocols were described previously (Zhao *et al*. 2020; Deanhardt *et al*. 2022).

Briefly, for each biological replicate, equal number of male and female heads (7 to 10, 5-7 days old post eclosion) were dissected in RNase-free environment with Trizol, followed by tissue homogenization, cell lysis, and filtration with Qiagen QIAshredder spin column. Three biological replicates were analyzed for each genotype. RNA was then extracted and purified by Qiagen RNeasy Kit per manufacturer’s instructions and eluted in 60 μL RNase-free water. Genomic DNA was then removed using Invitrogen TURBO DNA-free Kit per manufacturer’s instructions. RNA concentration was measured using NanoDrop after DNase treatment. Reverse transcription was performed by Invitrogen SuperScript IV First-Strand cDNA synthesis Reaction Kit per manufacturer’s instructions. Notably, approximately equal amount of template RNA across different samples were added based on RNA concentration.

Lastly, qPCR reactions were run on Roche LightCycler 96 Instrument with FastStart Essential DNA Green Master (2X) in 20 μL volume with technical triplicates per manufacturer’s instructions. Thermal cycling program: 600 s pre-incubation at 95 °C + 40 three-step amplification cycles (10 s denaturation at 95 °C, 10 s annealing at 55 °C, 15 s extension at 72 °C with Single acquisition) + Melting Curve (10 s 95 °C, 60 s 65 °C, 1 s 97 °C with 5 Readings/°C). qPCR primers were designed to span two adjacent exons when possible, with target amplicon length of 120 bp. Primer pairs were tested to generate standard curves to evaluate amplification efficiency before being used for expression comparison experiments. Expression levels of the gene of interest were normalized to the house keeping gene *RpL13A* in each sample by 2^-ΔCt^ method. Two-way ANOVA followed by multiple comparison test was performed in Prism 9 software to determine statistical significance.

**Table 1.**
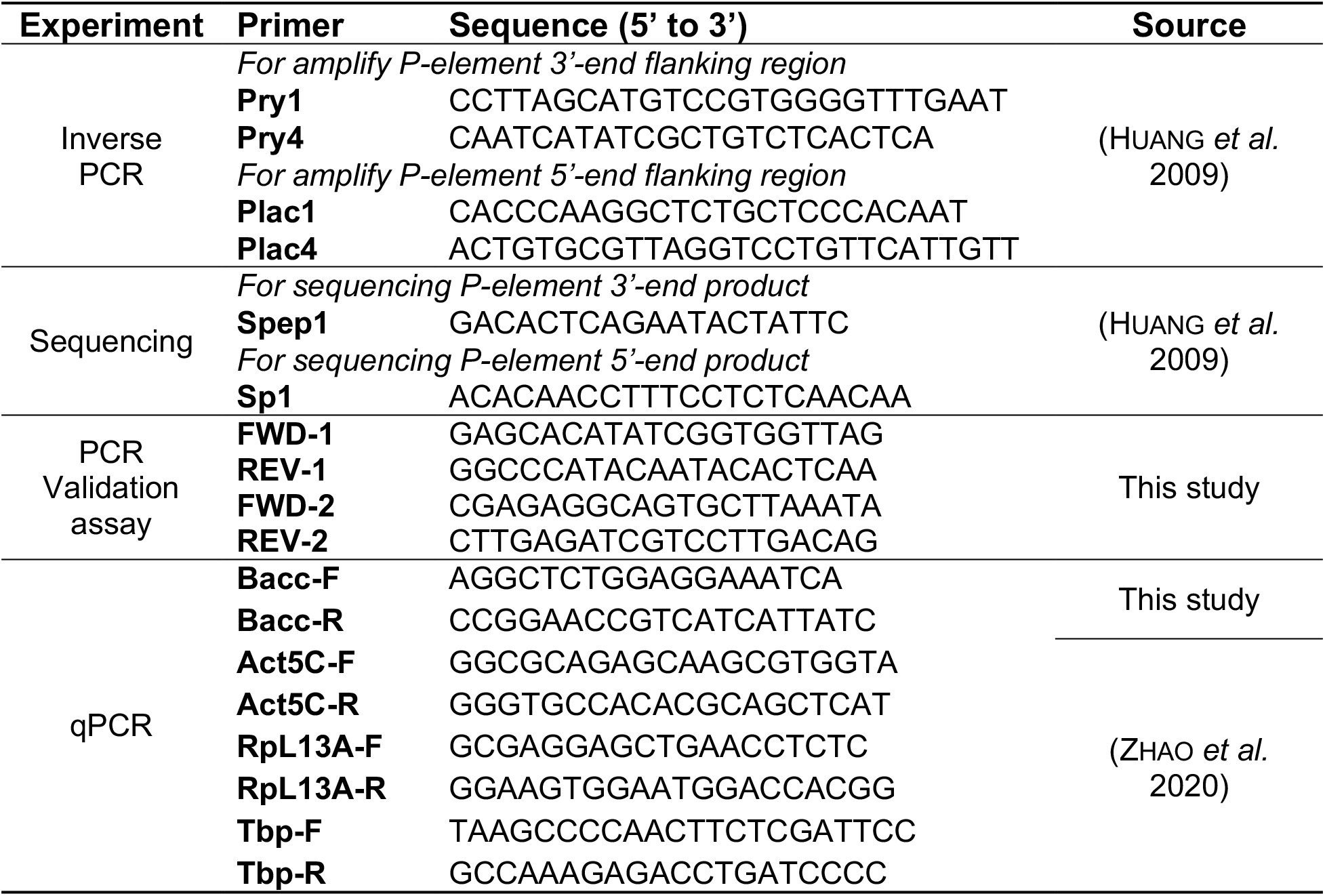
Primers used in this study.

## Data availability

The authors affirm that all the data necessary for drawing the conclusions are present in the text, figures, and figure legends. Most of the *Drosophila* stocks are obtained from Bloomington or Vienna stock center, with identifiers listed in the materials and methods section. All the other lines are available upon request.

## Acknowledgments

We would like to thank Bloomington Drosophila Stock Center and Vienna Drosophila Resource Center for providing all the fly stocks. We thank Duke Light Microscopy Core Facility for help with imaging. We thank Chengcheng Du for help with molecular biology.

## Funding

This study is funded by grant NSF 2006471 and NIH 5R01NS109401 (both to P.C.V.). Q.D. is supported by Duke Biology Department Ph.D. program.

## Conflicts of interest

The authors declare no conflict of interest with the contents of this paper.

## Author contribution

Q.D. and P.C.V. conceived the study and designed the experiments; Q.D. did most of the experiments with help from R.E., A.C., and Y.C.; Q.D. analyzed the data and prepared the figures; Q.D. and P.C.V. wrote and edited the manuscript.

---

**Figure S1.**
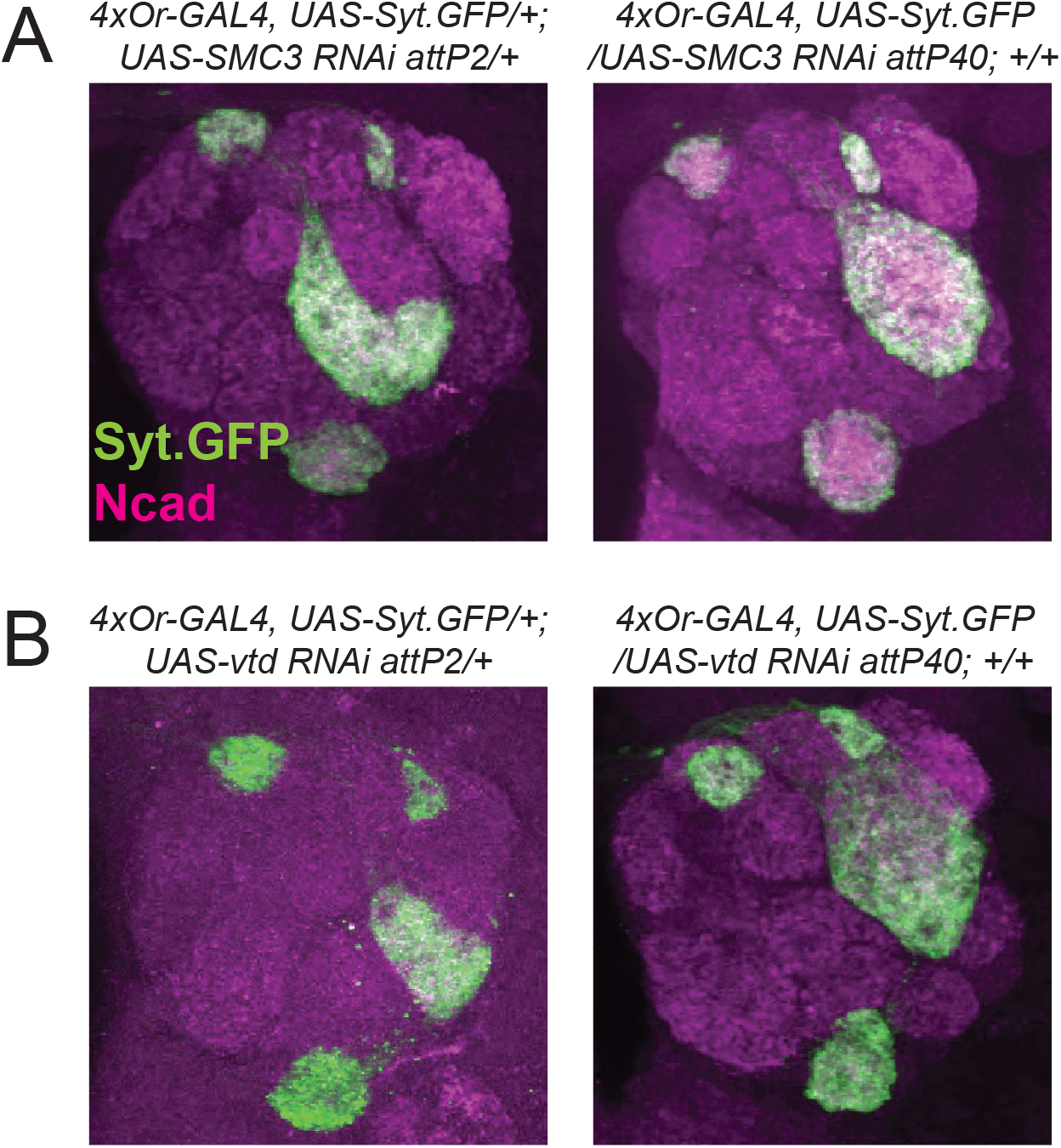
Additional *TRiP UAS-RNAi* results showing the *attP40* but not *attP2-* specific glomerular organization phenotype. Similar to *Beat/Side* screening results in Figure 1A,B, crossing *4xOr-GAL4, UAS-Syt*.*GFP* chromosome to two *UAS-SMC3 RNAi* lines **(A)** and *UAS-vtd RNAi* lines **(B)** respectively gave rise to glomerular expansion with *attP40* insertion but not *attP2* insertion. 5-9 brains were examined in each genotype and the phenotypical penetrance is 100% in *attP40* groups.

**Figure S2.**
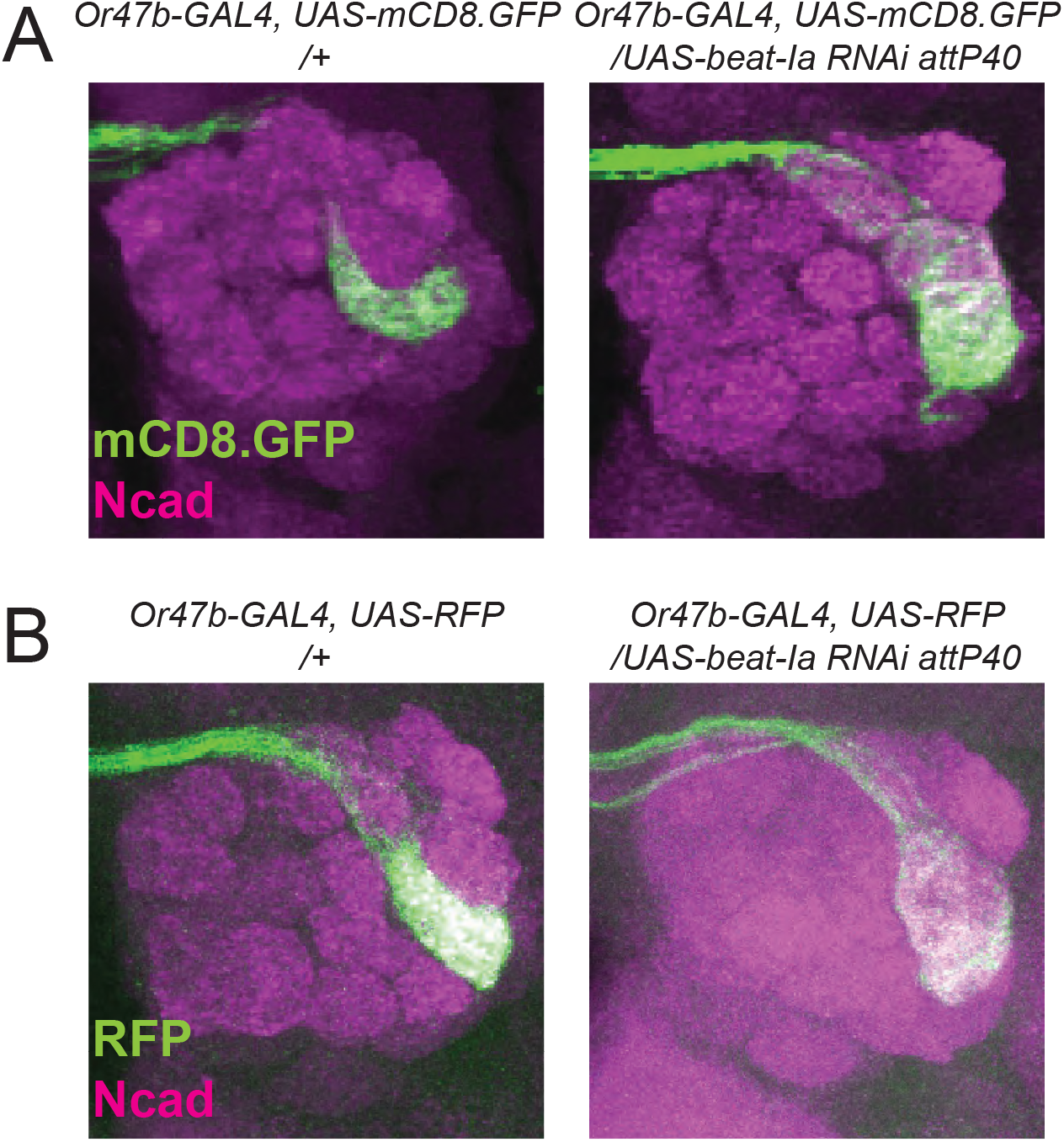
*attP40* over *Or47b-GAL4* recombined with *UAS-reporters* other than *UAS-Syt.GFP* exhibits the same glomerular organization phenotype. *Or47b-GAL4, UAS-mCD8*.*GFP* **(A)** and *Or47b-GAL4, UAS-RFP* **(B)** over the *attP40* derivative showed VA1v glomerular expansion compared with the respective no-*attP40* control. 5-15 brains were examined in each genotype and the phenotypical penetrance is 100% in *attP40* groups.

## Notes

### Competing Interest Statement

The authors have declared no competing interest.

### Summary of Updates

A new main figure (Figure 3) was added. The text was also modified accordingly. This new version is an accepted manuscript for publication.

